# pavo 2: new tools for the spectral and spatial analysis of colour in R

**DOI:** 10.1101/427658

**Authors:** Rafael Maia, Hugo Gruson, John A. Endler, Thomas E. White

## Abstract

1. Biological colouration presents a canvas for the study of ecological and evolutionary processes. Enduring interest in colour-based phenotypes has driven, and been driven by, improved techniques for quantifying colour patterns in ever-more relevant ways, yet the need for flexible, open frameworks for data processing and analysis persists.
2. Here we introduce `pavo 2`, the latest iteration of the R package `pavo`. This release represents the extensive refinement and expansion of existing methods, as well as a suite of new tools for the cohesive analysis of the spectral and (now) spatial structure of colour patterns and perception. At its core, the package retains a broad focus on (a) the organisation and processing of spectral and spatial data, and tools for the alternating (b) visualisation, and (c) analysis of data. Significantly, `pavo 2` introduces image-analysis capabilities, providing a cohesive workflow for the comprehensive analysis of colour patterns.
3. We demonstrate the utility of `pavo` with a brief example centred on mimicry in *Heliconius* butterflies. Drawing on visual modelling, adjacency, and boundary strength analyses, we show that the combined spectral (colour and luminance) and spatial (pattern element distribution and boundary salience) features of putative models and mimics are closely aligned.
4. `pavo 2` offers a flexible and reproducible environment for the analysis of colour, with renewed potential to assist researchers in answering fundamental questions in sensory ecology and evolution.

## Introduction

The study of colour in nature continues to generate fundamental knowledge: from the neurobiology and ecology of information processing (Caves *et al*., 2018; Schnaitmann *et al*., 2018; Thoen *et al*., 2014; White & Kemp, 2017), to the evolutionary drivers of life’s diversity (Dalrymple *et al*., 2015, 2018; Endler, 1980; Maia *et al*., 2013b). Colour is a subjective perceptual experience, however, so our understanding of the function and evolution of this conspicuous facet of variation depends on our ability to analyse phenotypes in meaningful ways. Excellent progress continues to be made in this area, with emerging techniques now able to quantify and integrate both the spectral (i.e. colour and luminance) and spatial (i.e. the distribution of pattern elements) properties of colour patterns (Endler, 2012; Endler *et al*., 2018; Kemp *et al*., 2015; Renoult *et al*., 2015; Troscianko *et al*., 2017). The need remains, however, for tools that integrate these complex methods into clear, open, and reproducible workflows (White *et al*., 2015), allowing researchers to retain focus on the exploration of interesting questions.

Here we introduce `pavo 2`, a major revision and update of the R package `pavo` (Maia *et al*., 2013a). Since its initial release, the package has provided a cohesive framework for the processing and analysis of spectral data, yet the interceding years have seen the advent of novel analytical methods and the refinement of existing ones. As detailed below, `pavo 2` has been extensively expanded to incorporate a suite of new tools, with the most significant advance being the inclusion of geometry-based analyses. This allows for the quantification of spectral and spatial properties of colour patterns within a single workflow, thereby minimising the computational and cognitive overhead associated with their otherwise fragmented analysis.

## The pavo package, version 2

The conceptual focus of pavo remains centred on three components: (1) data importing and processing, and ongoing feedback between (2) visualisation and (3) analysis (Fig. 1). The package is available for direct installation through R from CRAN (https://CRAN.R-project.org/package=pavo), while the development version remains available on Github (https://github.com/rmaia/pavo). Comprehensive details and examples of the rich functionality of `pavo` are available in help files as well as the package vignettes. Indeed, we strongly encourage readers to refer to the vignettes as the primary source for information on `pavo`’s functionality (accessible `through browseVignettes(pavo)`, and at http://rafaelmaia.net/pavo/), since they are updated as necessary with every package release.

**Figure 1:**
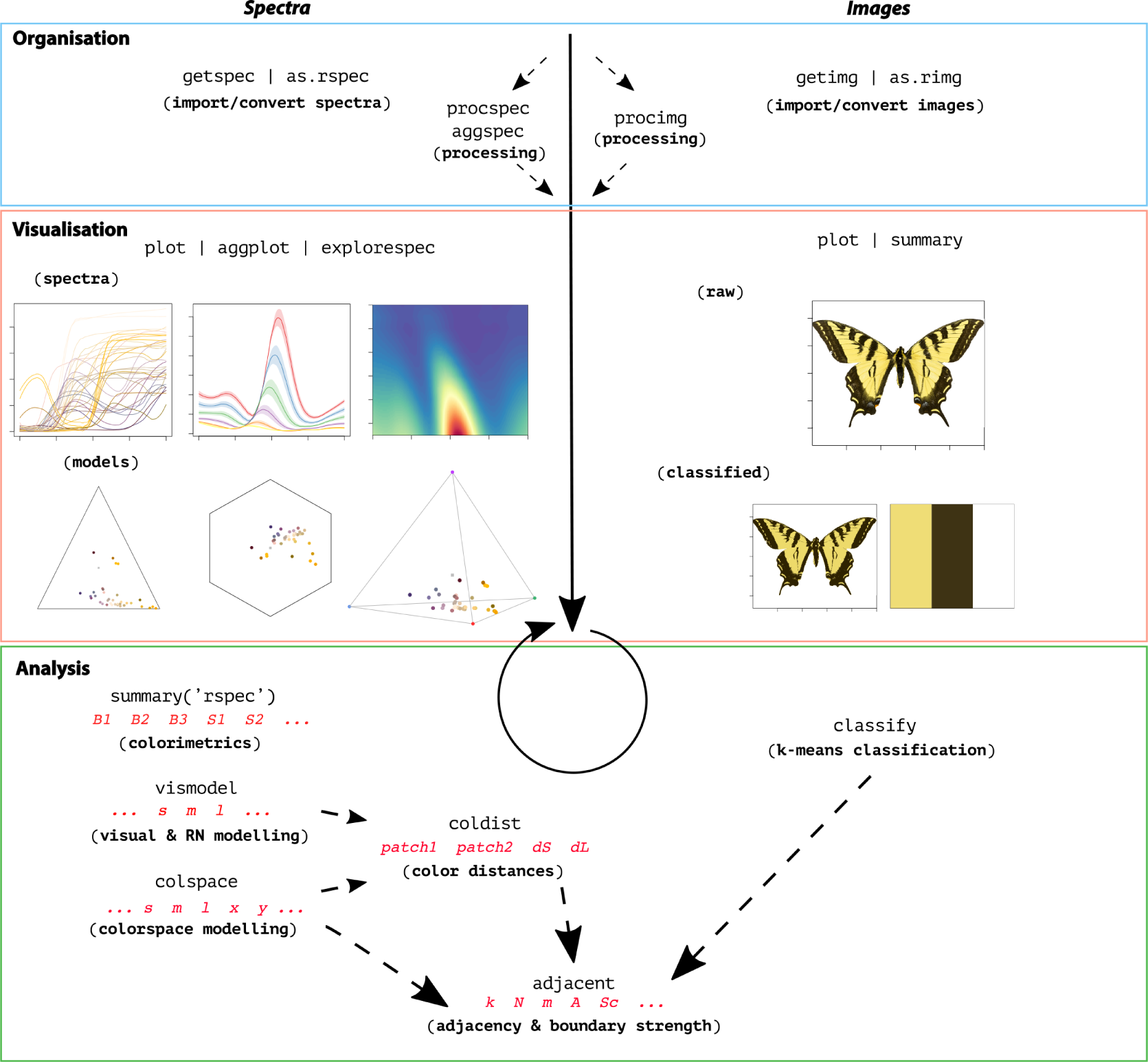
A general overview of the colour-pattern analysis workflow in pavo, as of version 2, displaying some key functions at each stage.

### Organisation

Images and spectra can be loaded into `pavo` in bulk through the use of `getimg()` and `getspec()`, respectively. Both are capable of handling multiple data formats, such as jpeg, bmp and png in the case of images, and over a dozen formats of spectral data, including the diverse and complex proprietary formats of the various spectrometer vendors. Once loaded, the data are stored as objects of an appropriate custom S3 class, for use in further functions. Spectral data are of class `rspec`, and inherit methods from `data.frame`, while images are of class getimg, and are multidimensional objects (typically 3D, for an RGB image) that inherits methods from `array`. If more than one image is imported in a single call to `getimg()`, then each image is stored as an element of a `list`. This class system allows for — among other things — the reliable use of generic functions such as `plot()` and `summary()`, which can be called any time to inspect and visualise data.

Several functions then facilitate the initial processing of colour data. It is often desirable to process spectra to remove unwanted noise, modify the spectral range, and/or interpolate the standard wavelength intervals, all of which may be achieved through `procspec()`. For images, `procimg()` offers similar functionality such as the ability to interactively specify the real-world scale of images (in preferred units of measurement), rotate and resize images, or define the boundary between a focal object and the visual background. The scope of image processing in `pavo 2` is relatively limited by design, as much of what might be used during standard image handling are either needs best considered and met by researchers during image capture and data-checking, or are readily achieved within R using existing packages such as `imager` (Barthelme, 2018) and `magick` (Ooms, 2018). Indeed, `pavo 2` includes convenience functions to convert between image-classes used by `pavo`, `imager`, and `magick`, allowing ready access to extensive image-processing capabilities.

### Visualisation

The repeated visualisation of spectral and spatial data is an essential step during all stages of analysis, and `pavo 2` offers numerous tools and publication-ready graphics fit for purpose. Once the package is loaded, the `plot()` function recognises objects of class `rspec` and `rimg`, as well as `colspace` (the product of visual modelling, detailed below), and becomes the conduit to most visualisations. For raw spectral data, for example, `plot()` will produce a clean plot of the spectra versus wavelengths (Fig. 1, centre-left). Following visual modelling, di-, tri-, and tetra-chromatic models can instead be visualised, as well as data from more specialised models, such as the colour hexagon (Chittka, 1992), CIEXYZ or LAB spaces (Smith & Guild, 1931; Westland *et al*., 2012), categorical space (Troje, 1993), segment analysis (Endler, 1990), the colour-opponent coding space (Backhaus, 1991), or the ‘receptor-noise’ space (de Ibarra *et al*., 2001; Pike, 2012). Images can also be plotted, with the result depending on whether and how they have been processed. When given an unprocessed `rimg` object, `plot()` will produce a simple raster-based plot of the image (Fig. 1, right). Following the results of `classify()` (discussed below), in which image pixels are k-means classified into discrete colour-classes (or if a colour-classified image is loaded directly), the plot will use the mean RGB values of each colour-class to plot the now-classified image (Fig. 2).

**Figure 2:**
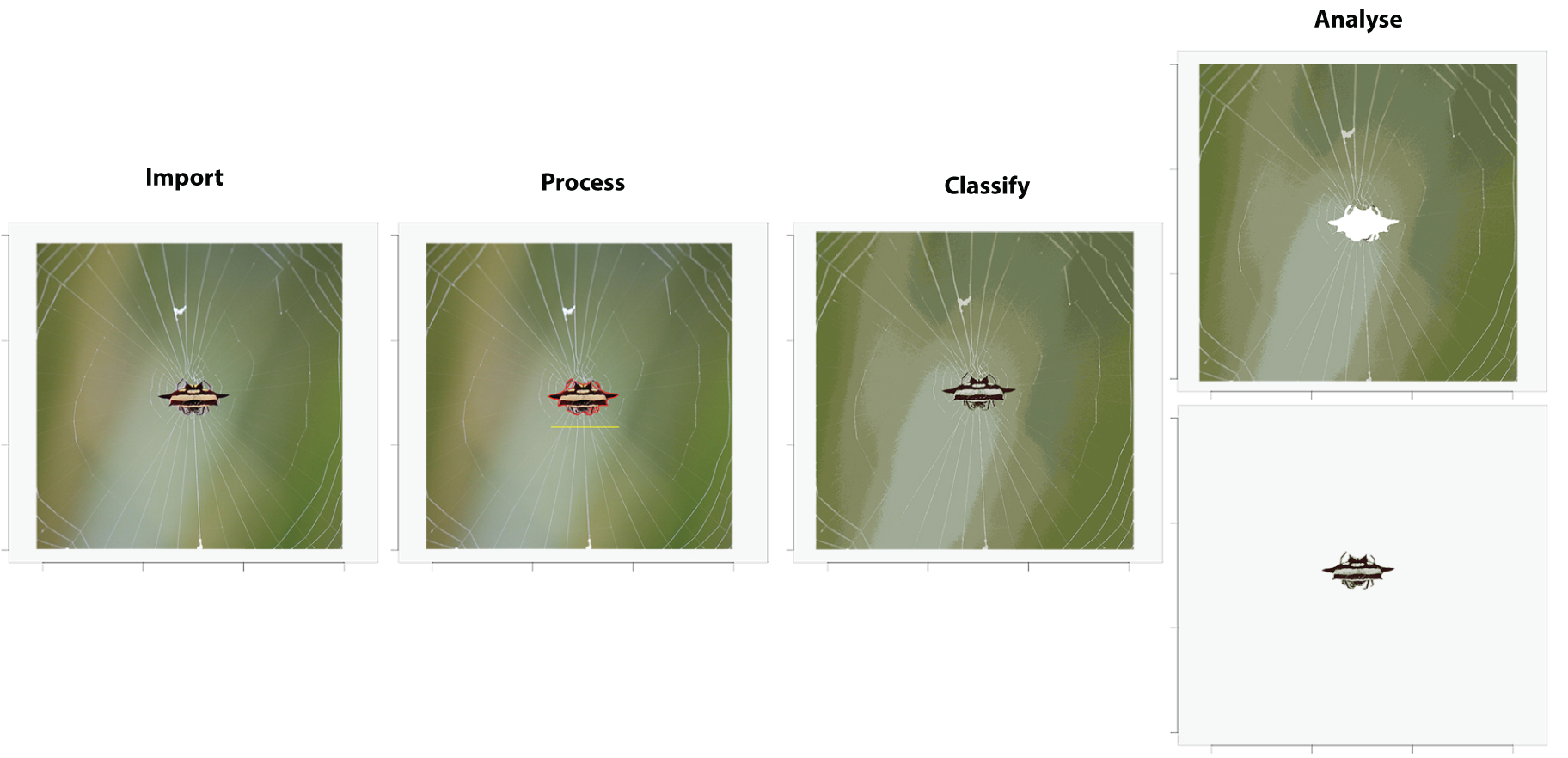
A sample workflow for image handling and analysis in pavo, as of version 2. Images are first imported and optionally processed by, for example, setting scales (yellow line) or defining objects and backgrounds (red outline). They may then be colour-classified before being passed to analytical functions, currently centered on the adjacency and boundary-strength analyses. If backgrounds and focal objects are defined then they can be analysed separately, concurrently, or either one can be excluded entirely.

### Analysis

Since the perception of colour is a subjective experience, significant progress has been made in representing its reception using ecologically relevant ‘visual models’ (Kelber *et al*., 2003; Kemp *et al*., 2015; Renoult *et al*., 2015), which `pavo 2` includes in an extended repertoire. The first step in such analyses is a call to `vismodel()`, which models photoreceptor stimulation (quantum-catches, or photon-flux) based on information about the viewer’s visual sensitivity and viewing environments. While users are free to use their own spectra, `pavo` includes a suite of built-in receptor sensitivities, illuminant and transmission data (be it environmental or ocular), and viewing backgrounds, for convenience.

Once quantum catches are estimated the results can used in a number of models, depending on the question and analytical objective at hand (Kemp *et al*., 2015; Renoult *et al*., 2015). General colourspaces are available through a call to `colspace()` which, if provided no further arguments, will model the data in a generalist ditri- or tetrachromatic space informed by the dimensionality of the visual system. More specialised colourspaces — which may be informed by specific information about the visual perception of particular species — are also available via `colspace()`. The CIEXYZ, CIELAB, and CIELch models (designed and intended exclusively for humans) are available, and `colspace()` will check that the appropriate inputs, such as the human colour-matching function, have been used to model receptor stimulation, as required (Smith & Guild, 1931; Westland *et al*., 2012). The colour-opponent-coding (Backhaus, 1991) and colour-hexagon (Chittka, 1992) models of bee vision are implemented, as is the categorical model of fly colour-vision detailed by Troje (1993). Plots for every space are accessible through a call to `plot()` which, thanks to the underlying class system, will draw on the appropriate visualisation for the model at hand — be it a hexagon, a dichromatic segment, a Maxwell triangle, or a three-dimensional tetrahedron.

The receptor-noise limited model of early-stage (retinal) colour processing has proven exceptionally popular (Vorobyev *et al*., 2001; Vorobyev & Osorio, 1998), and has been tested to varying degrees in diverse taxa (Barry *et al*., 2015; Fleishman *et al*., 2016; Kelber *et al*., 2003; Olsson *et al*., 2015; White & Kemp, 2016). Following the estimation of receptor stimulation in `vismodel()`, the model incorporates information on relative receptor densities and noise through the function `coldist()`, and estimates either quantum- or neural-noise weighted colour distances. Version 2 of `pavo` introduces several extensions of this approach, such as the bootstrapped colour distance of `bootcoldist()`, which provides an estimate of the noise-weighted distances (*δS*’s and/or *δL*’s) between the centroids of colour samples in multivariate space, with an appropriate measure of error (detailed in Maia & White, 2018). Stimuli can also now be expressed and plotted as coordinates in ‘perceptual’ (i.e. receptor-noise corrected) space by calling `jnd2xyz()` on the distances calculated in `coldist()` (de Ibarra *et al*., 2001; Pike, 2012). Notably, these functions now accept n-dimensional data (derived independently, but see Clark *et al*., 2017; Gawryszewski, 2018, for valuable discussion). This allows for the modelling of extreme (Chen *et al*., 2016; Cronin & Marshall, 1989, though given the lack of support for traditional opponency in these systems, the RN model may be of limited use here) or entirely hypothetical visual systems. Of course `coldist()` also accepts the results of alternative models — such as the hexagon or CIELab — and will return colour distances in units appropriate for each space.

Exciting recent advances now allow for the analysis of colour pattern geometry — that is, the *spatial* structure of colour patches — in conjunction with the comparatively well-developed approaches to the *spectral* analysis of colour out-lined above (Endler, 2012; Endler *et al*., 2018; Pike, 2018; Troscianko *et al*., 2017). The most significant extension of `pavo` as of version 2 is the introduction of an image-based workflow to allow for the combined analysis of the spectral and spatial structure of colour patterns, currently centred on measures of overall pattern contrast (Endler & Mielke, 2005), the adjacency analysis (Endler, 2012), and its extension, the boundary strength analysis (Endler *et al*., 2018). In `pavo 2`, the various steps for such analyses are carried out through calls to `classify()`, which uses k-means clustering to automatically or interactively classify image pixels into discrete colour-classes, and/or `adjacent()`, which performs the adjacency analysis and, if appropriate colour distances are also specified, the boundary strength analysis (discussed in Endler *et al*., 2018).

Briefly, these analyses entail classifying evenly-spaced points within a visual scene into discrete colour classes using spectrometric measurements and/or photography. The column-wise and row-wise colour-class transitions between adjacent points are then tallied, and from this a suite of summary statistics on pattern structure — from simple colour proportions, through to colour diversity and pattern complexity — are estimated (e.g. Endler *et al*., 2014; Rojas *et al*., 2014; Rojas & Endler, 2013; White, 2017). The precise procedure that might be followed by researchers may vary considerably depending on the goal and tools at hand, and `pavo 2` is designed to accommodate such flexibility. In relatively simple cases (as in the below example), users may import and calibrate images via `getimg()` and `procimg()`, k-means classify the entire image using `classify()`, and combine it with spectrometric measurements and visual modelling of the few discrete colour-classes in a call to `adjacent()`. In more complex cases, such as animals in their natural habitats, users may instead wish to collect spectrometric measurements along a grid-sample of the visual scene, visually model and statistically cluster the results (e.g. using `vismodel()`), then feed the resulting colour-classified grid into `adjacent()` directly (as per ‘method 1’: Endler, 2012), without the use of images or the `classify()` function at all.

As alluded to earlier, our goal is to provide a flexible and relatively simple analytical framework for the analysis of a colour pattern’s spatial structure using images, without the requirement for specialised photographic equipment or and/or extensive calibration and processing (demonstrated in the colour-plate based example below). We thus make an analytical and conceptual distinction between the spectral data afforded by spectrometry, and the spatial data afforded by images, with the two able to be conveniently combined during latter analyses (Fig. 1). This also minimises the unnecessary duplication of efforts of more general-purpose tools such as imager (Barthelme, 2018) and `magick` (Ooms, 2018), and the excellent image analysis toolbox for imageJ (Troscianko & Stevens, 2015), which offer rich functionality for image processing and (in the latter case) analysis. We emphasise, however, that the convenience of the toolkit provided by `pavo 2` belies the complexity of the choices demanded of researchers, and that every parameter and option requires close consideration and justification. It is rare, for example, that image analyses should be used without any input from visually-modelled spectrometric data, since naive clustering performed on uncalibrated images will typically offer a poor representation of a visual scene as relevant to non-human animals. For example, even in simple cases, as below, the number of discrete patches present (i.e. the argument `kcols` in `cluster()`) is best estimated using spectro-metric data in an ecologically relevant model, rather than relying exclusively on human-subjective estimates of colour segregation. One possible approach is integrated into the below example, and Endler (2012) details others, such as estimating `kcols` as the number of receptor-noise ellipsoids required to encompass the entire sample of spectra.

## Worked example: mimicry in *Heliconius* spp

Butterflies of the genus *Heliconius* are widely involved in mimicry, and have proven an exemplary system for studies of colour pattern development, ecology, and evolution (Jiggins, 2016). Here we demonstrate some of `pavo 2`’s capabilities by briefly examining the the visual basis of mimicry in this system, with the objective of quantifying the spectral and spatial (dis)similarity between putative models and mimics. For our spatial analyses, we follow Endler (2012) and use colour plate XII from Eltringham (1916), which is arranged into what he described as model and mimic pairs (Fig. 3). For our spectral analyses we collated six reflectance spectra from each of the assumed-discrete ‘red’, ‘yellow’, and ‘black’ patches (confirmed by spectral measurement, below) of the forewings of two species — *H*. *egeria* and *H*. *melpomene* (Fig. 3, top left pair) — from personal sources and the literature (Bybee *et al*., 2011; Wilts *et al*., 2017). For reasons of simplicity and data availability we restrict our visual modelling to these two species, though the below spectral analyses would ideally be repeated for all model/mimic pairs.

**Figure 3:**
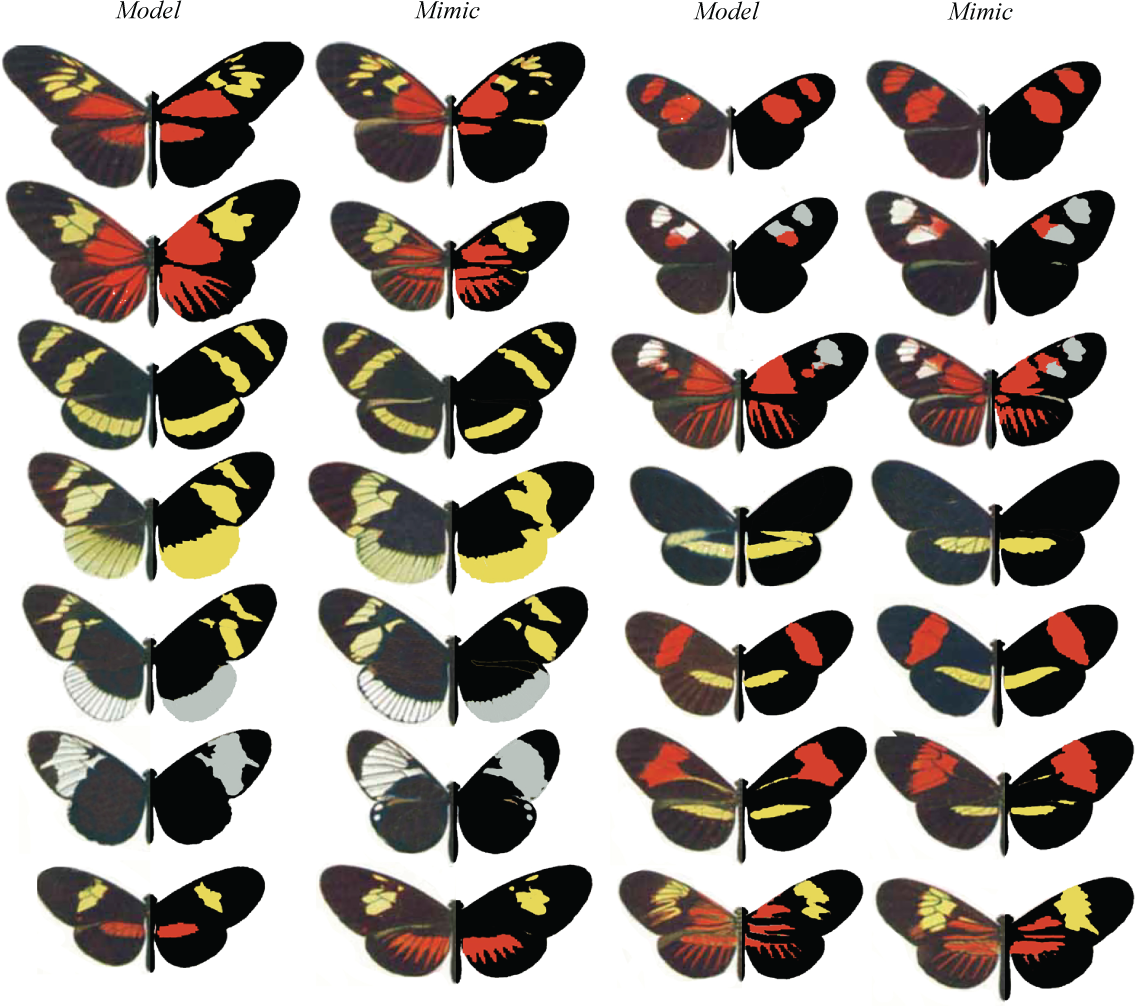
A modification of Eltringham’s (1916) colour plate of *Heliconius* butterflies, *sensu* Endler (2012), arranged into putative models and mimics. The left side of each individual is as per the original, while the right half display pattern elements that have been classified into discrete classes through k-means clustering, using the `classify()` function.

### Spectral analysis

We first focus on the spectral data, both to confirm the assumption that there are discrete colour patches and because some of the results of this work will be drawn on for the latter pattern analyses. We begin by loading the reflectance spectra, which are saved in a single tab-delimited text file along with the image plates (available at the package repository; https://github.com/rmaia/pavo, or via figshare; https://dx.doi.org/10.6084/m9.figshare.7445840.v1), before LOESS-smoothing them to remove any minor electrical noise and zeroing spurious negative values.

~~~
*# Load spectra*

> heli_specs <-getspec('../data', ext = 'txt')

*# Smooth spectra and zero negative values*

> heli_specs <-procspec(heli_specs,

>                       opt = 'smooth',

>                       fixneg = 'zero')
~~~

A call to plot(`heli_specs, col = spec2rgb(heli_specs`)) displays the now-clean spectra, with each line coloured according to how it might appear to a human viewer (Fig. 4, top left).

**Figure 4:**
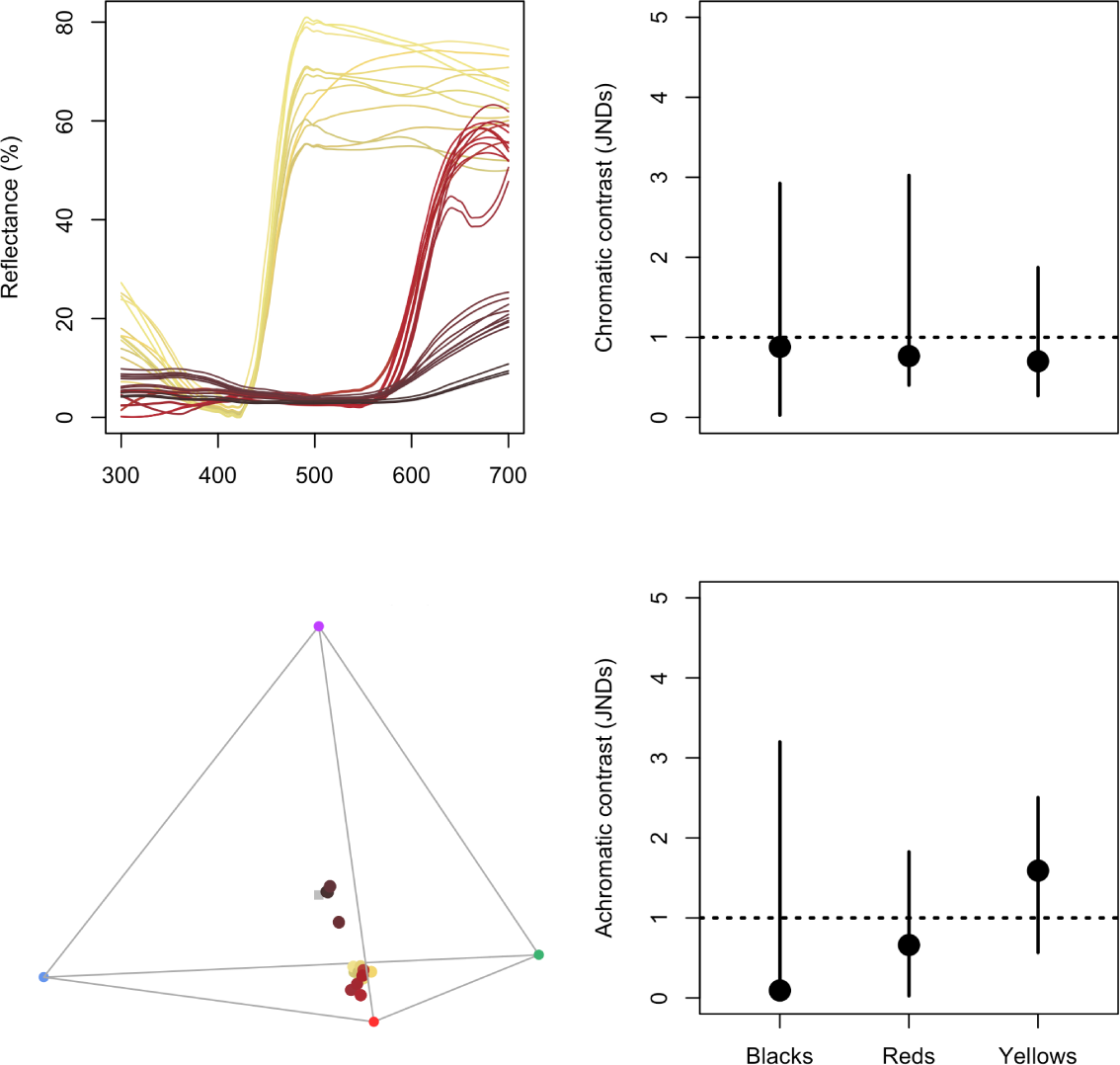
Reflectance spectra from black, red, and yellow patches of *H*. *egeria* and *H*. *melpomene*, along with their positions in a tetrahedral model of avian vision (left side). The bootstrapped, noise-corrected chromatic and achromatic patch distances between species (right) predicts that the individual colours of this model/mimic pair are likely indistinguishable to avian predators.

Our interest is in quantifying the fidelity of visual mimicry, so we must consider the perspective of ecologically relevant viewers (the primary selective agents) which, in the case of aposematic *Heliconius*, are avian predators (Benson, 1972; Chai, 1986). We thus use the receptor-noise limited model (Vorobyev *et al*., 2001; Vorobyev & Osorio, 1998) to predict whether the black, red, and yellow colour patches of a representative model and mimic are distinguishable to avian predators. This first entails estimating the photoreceptor quantum catches of a repre-sentative viewer, so we use a built-in average UV-sensitive avian visual phenotype for estimating chromatic distances, and the double-cone sensitivity of the blue tit for luminance distances.

~~~
> heli_model <-vismodel(heli_specs,

>                       visual = 'avg.uv',

>                       achromatic = 'bt.dc',

>                       relative = FALSE)
~~~

At this point we may wish to get a quick sense of the relative distribution of stimuli by converting them to locations in an avian tetrahedral colourspace and plotting the results with plot(`colspace(heli_model)`) (Fig. 4). With receptor stimulation estimated, we now calculate noise-corrected chromatic and achromatic distances between patches. The `coldist()` function can be used to return the pair-wise distances between every spectrum, which might then be averaged to derive a mean distance between species for every patch. This neglects the multivariate structure of such data, however, when the objective is to estimate the separation of groups in colourspace (Maia & White, 2018). We therefore prefer a bootstrapped measure of colour distance using `bootcoldist()`, which provides a robust measure of the separation of our focal samples (i.e the red, white, and black patches of model versus mimic), along with a 95% confidence interval, which can be inspected to see if it exceeds the theoretical discrimination threshold of one JND. We specify a relative receptor density of 1:2:2:4 (ultraviolet:short:medium:long wave-length receptors; Maier & Bowmaker (1993)), a signal-to-noise ratio yielding a Weber fraction of 0.1 for both chromatic and achromatic receptors, and assume that noise is proportional to the Weber fraction and independent of the magnitude of receptor stimulation (reviewed in Kelber *et al*. (2003); Olsson *et al*. (2017)).

~~~
*# Calculate the bootstrapped, noise-corrected colour distance*

*# between groups, using sample names to specify grouping ID’s*.

> heli_dist <-bootcoldist(heli_model,

>                         by = sub('\\..*', '', rownames(heli_model)),

>                         n = c(1, 2, 2, 4),

>                         weber = 0.1,

>                         weber.achro = 0.1)
~~~

Inspection of the key comparisons of interest (Fig. 4, right) reveals that the 95% CI of all chromatic and achromatic comparisons includes the theoretical threshold of one JND. This predicts that the individual colour pattern elements of putative model and mimic *H*. *egeria* and *H*. *melpomene* are indistinguishable, or difficult to discriminate, to avian viewers — the assumed intended recipient of the aposematic signals. As noted above, the analysis of this representative pair can be readily scaled to encompass all species given the necessary data, and we can now use this information to inform our study of the spatial structure of these signals.

### Pattern analysis

We first load the focal images, which comprise the individual samples from plate XII of Eltringham (1916), saved as jpegs (Fig. 3). We then plot one or all of the images to check they are as expected.

~~~
*# Load all images*. *Here the 28 jpegs are stored in a folder called*

*# 'butterflies' located within the current working directory*.

> heli_images <-getimg(“butterflies”)

28 files found; importing images.

*# Plot the first image in the list only*.

> plot(heli_images[[1]])

*# Plot all images, which will progress through

# the sequence automatically*.

> plot(heli_images)
~~~

We then classify the pixels of all images into discrete colour or luminance categories, here using k-means clustering, to create a colour-classified image matrix. The function `classify()` will carry this out, though there are numerous specific ways in which it may be achieved, including automatically or ‘interactively’, with the option of a reference image as template. Since our images are heterogeneous, it is simplest to use the interactive version of `classify()`, which will cycle through each image and ask the user to manually identify a sample from every discrete colour or luminance class present, which are then used as cluster centres.

~~~
*# Interactively colour-classify all images using k-means clustering*.

> heli_class <- classify(heli_images, interactive = TRUE)

*# Cycle through plots of the colour-classified images, alongside their

# identified colour palettes*.

> summary(heli_class, plot = TRUE)
~~~

Finally, we use an adjacency analysis to estimate a suite of metrics describing the structure and complexity of the colour pattern geometry of model and mimic *Heliconius*, and by including the visually-modelled colour distances estimated above, the output will include several measures of the salience of colour patch edges as part of the boundary strength analysis (Endler, 2012; Endler *et al*., 2018). We will exclude the white background since it is not relevant, simply by specifying the colour-category ID belonging to the homogeneous underlay. If the image was more complex, such as an animal in its natural habitat, we might instead interactively identify and separate the focal animal and background using `procimg()` (e.g. Fig. 2, second panel). Alternatively, we might forego the use of images altogether, and instead grid-sample and cluster the spectra across the visual scene and use these in directly in the call to `adjacent()` (*sensu* ‘method 1’ in Endler 2012, mentioned above).

~~~
*# Construct and inspect a data.frame of pairwise colour and luminance*

*# distances between all colour classes, built from the earlier*

*# receptor-noise modelled estimates. Note that we do not bother*

*# including colour-class ID 1, since that is the white background*

*# which is to be excluded from the analysis (see below)*.

*# (Alternatively we could include it, and it would simply be ignored)*.

> distances <-data.frame(c1 = c(2, 2, 3),

                         c2 = c(3, 4, 4),

                         dS = c(10.6, 5.1, 4.4),

                         dL = c(1.1, 2.5, 3.2))
> distances

c1  c2  dS     dL

2   3   10.50  7.41

2   4   11.76  23.40

3   4   13.29  15.99

*# Calculate adjacency and boundary-strength statistics. We specify a*

*# scale of 50 mm, and note that the 'white' background, which has the class*

*# ID of 1 in this case, is to be excluded from the analysis*.

*# We also include the colour distance between all patches, as estimated above*.

> heli_adj <-adjacent(heli_class,

>                     xscale = 50,

>                     bkgID = 1,

>                     exclude = 'background',

>                     coldists = distances)

*# Inspect a subset of the resulting data.frame. Variable meanings*

*# are detailed in the function documentation (see ?adjacent)*,

*# or Endler (2012), Endler et al. (2018), and Endler & Mielke (2005)*.

> head(heli_adj)[, 1:7]

          k  N        n_off   p_2    p_3    p_4    q_2_2 …

 mimic_01 3  345522   6547    0.801  0.130  0.067  0.796

 mimic_02 2  1018370  4091    0.835  0.164  NA     0.834

 mimic_03 3  265278   6155    0.685  0.198  0.116  0.677

…
~~~

We can now inspect the pattern descriptors of particular interest, and explore the similarity of models and mimics with respect to their broader colour pattern geometry. As seen in Fig. 5, the relative proportions of focal colours (top row), measures of pattern diversity and complexity (centre row), and the salience of patch boundaries (bottom row) are highly correlated between species pairs. This, in conjunction with the above modelling, suggests that the overall colour patterns of putative model and mimic *Heliconius* — both spectrally and spatially — are highly similar, and are thus predicted to be very difficult to discriminate to the intended avian viewers of their aposematic signals, as consistent with theory (Müller, 1879). More interesting questions remain, of course, including the degree to which mimics need resemble models to deceive viewers, and the relative importance of different colour pattern elements (e.g. Fig. 5) in mediating the subjective resemblance of species pairs, for which `pavo 2` is well suited to help answer.

**Figure 5:**
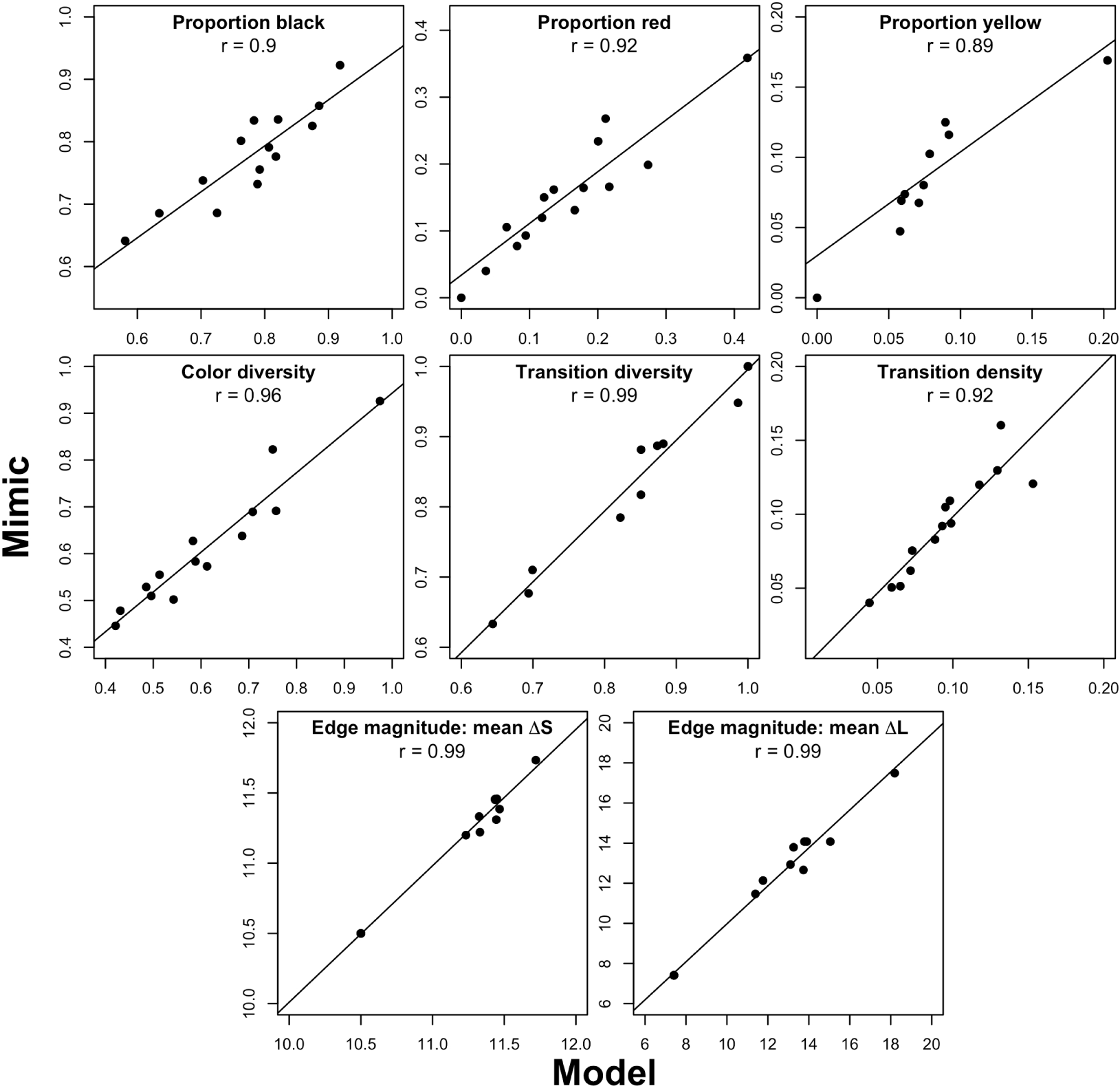
Select results of the colour pattern analysis of model and mimic *Heliconius* (Fig. 3), using adjacency and boundary strength analyses. Strong correlations are evident in colour proportions (top row), measures of colour diversity and complexity (centre row), and estimates of mean chromatic and achromatic edge salience (bottom row).

## Conclusions

The integrative study of biological colouration has borne rich fruit, though its potential to illuminate the structure and function of much of the natural world is not nearly realised (Endler & Mappes, 2017). As we have sought to demonstrate, `pavo 2` (and beyond) provides a flexible framework to assist researchers studying the physiology, ecology, and evolution of colour patterns and visual perception. We appreciate bug reports and suggestions, via email or the Github issue tracker https://github.com/rmaia/pavo/issues.

## Citation of methods

Many of the methods applied in `pavo 2` are described in detail in their original publications — as listed in the documentation for the relevant functions — to which users should refer and cite as appropriate, along with `pavo` itself, via this publication.

## Acknowledgements

We thank Kate Umbers, Georgia Binns, and Julia Riley for the rigorous testing of image-based methods. The package and manuscript also greatly benefited from the thoughtful input of an associate editor and three reviewers at MEE, which we appreciate. TEW thanks Elizabeth Mulvenna and Cormac White for their support. The authors have no conflicts of interest to declare.

## Authors statement

TEW, RM, and HG authored the software and manuscript, JAE developed and assisted in the implementation of methods, and critically revised the manuscript.

